# The MooSciTIC project: a booster shot of science for West African research and higher education actors

**DOI:** 10.1101/504092

**Authors:** Ménonvè Atindehou, Kifouli Adéoti, Laura Estelle Yêyinou Loko, Thierry Beulé, Emmanuel Paradis, Gustave Djedatin, Christine Tranchant-Dubreuil, François Sabot, Latifou Lagnika, Estelle Jaligot

## Abstract

In this paper, we describe the implementation of the MooSciTIC summer schools, a capacity-building initiative targeting West African research scientists and higher education teachers. Throughout the three years of the project, we provided on-site courses and practicals on cross-disciplinary aspects such as scientific writing, communication and integrity, in the aim of improving the self-reliance of researchers and upgrading research practices throughout the sub-region. We explain how the program was designed and implemented and we show the positive outcomes it generated for our former trainees within a relatively short period, hoping to inspire similar initiatives to follow our steps.

## Introduction

Over the past decades, a variety of short scientific training programs targeting developing countries have emerged. Regardless of the teaching approach used (on-site or distance learning) or the topics included in these programs, they share high requirements in human and infrastructural resources and thus heavily rely on external funding from developed countries or international organizations in order to access these resources and secure them over time. As a result, repeated occurrences of these trainings make them unsustainable to maintain over time because of the scarcity and competitiveness of funding sources, whereas the average life-span of a project (2-3 years) is generally regarded as insufficient to achieve the long-term, large-scale impact that is expected from such programs. There is no denying that the various teaching initiatives targeting developing countries have played a pivotal role in the curriculum of dozens of graduate students over the years, some of whom have since achieved tenure in their home institutions or abroad. Nevertheless, their impact is very small in comparison with the several thousands of undergraduate students attending African Universities each year in each discipline. Across the continent, it has indeed been noted that the number of new higher education institutions increases steadily to accommodate the demography but that this trend is not paralleled by a commensurate increase in government investments [1–3].

During our own previous teaching experiences across Western Africa, we had the opportunity to assess some of the specific difficulties faced by young research scientists and teaching assistants within this sub-region. In particular, we observed that their training generally did not include proper courses on key research-oriented aspects such as literature search, article and research proposal writing, project management, scientific integrity and ethics. Such knowledge gaps have been pointed out in other studies, often by African scientists themselves [4,5]. In high-income countries, such cross-disciplinary notions are commonly provided at postgraduate levels through a combination of courses and mentoring by senior scientists during lab internships. Thus, students from the North are progressively involved in research projects and the writing of publications and start building up their professional network, which strongly contributes to their early exposure to and integration within the currently highly competitive research environment, and ultimately, to their impact as scientists. By contrast, the African research landscape is complex and highly fragmented and, especially in French-speaking countries, it suffers from a lack of a culture of collaborative work and networking, including at the intra-national and regional levels. It can be assumed that such a deficit in networking may make African scientists less responsive to opportunities and their research less likely to achieve significant impact [6–8]. Sub-Saharan Africa accounts for an extremely small fraction of the world’s scientific publications (<1%, excluding South Africa), and it is even less represented among peer-reviewed ones [3,9,10]. In articles that are co-authored by scientists from the North and from the South, it is unfrequent to find the latter in last/senior author and corresponding author positions [11]. As scientists from developing countries, African scientists are also preferential preys to the so-called “predatory publishers” due to a disproportionate pressure towards quantity over quality of publications and their unfamiliarity with the current landscape of scientific publishing. As a result, their research further fails to gain international recognition [12,13]. As for international competitive grants, which are generally funded by North-based entities, West African scientists appear mostly as partners of the North-based institutions leading the project and they are not always part of the decision-making process. Under the combined effect of their lack of scientific visibility associated to insufficient publication record in legitimate, qualitative outlets, and their lack of practice in project development and management, they almost never assume coordinating positions. Self-evidently in these matters, lacking experience often leads to both being less likely to be offered opportunities, and not feeling confident enough to seize them. This constitutes a vicious circle (Fig 1) reminiscent of the well-described Matthew effect [14] that makes it harder for West African scientists to achieve international recognition for themselves. This phenomenon *de facto* places them in a relationship of strong, continued dependency towards their partners from high-income countries for the access to high-quality publications and competitive funding and, further down the road, the results do not always benefit research priorities of the South [5,7,8,15,16]. Ultimately, these constraints to their research greatly hampers the ability of West African scientists to teach their students by example how to address the same challenges they face themselves, namely getting their work funded, published and recognized beyond their borders by the greater scientific community.

**Fig 1.**
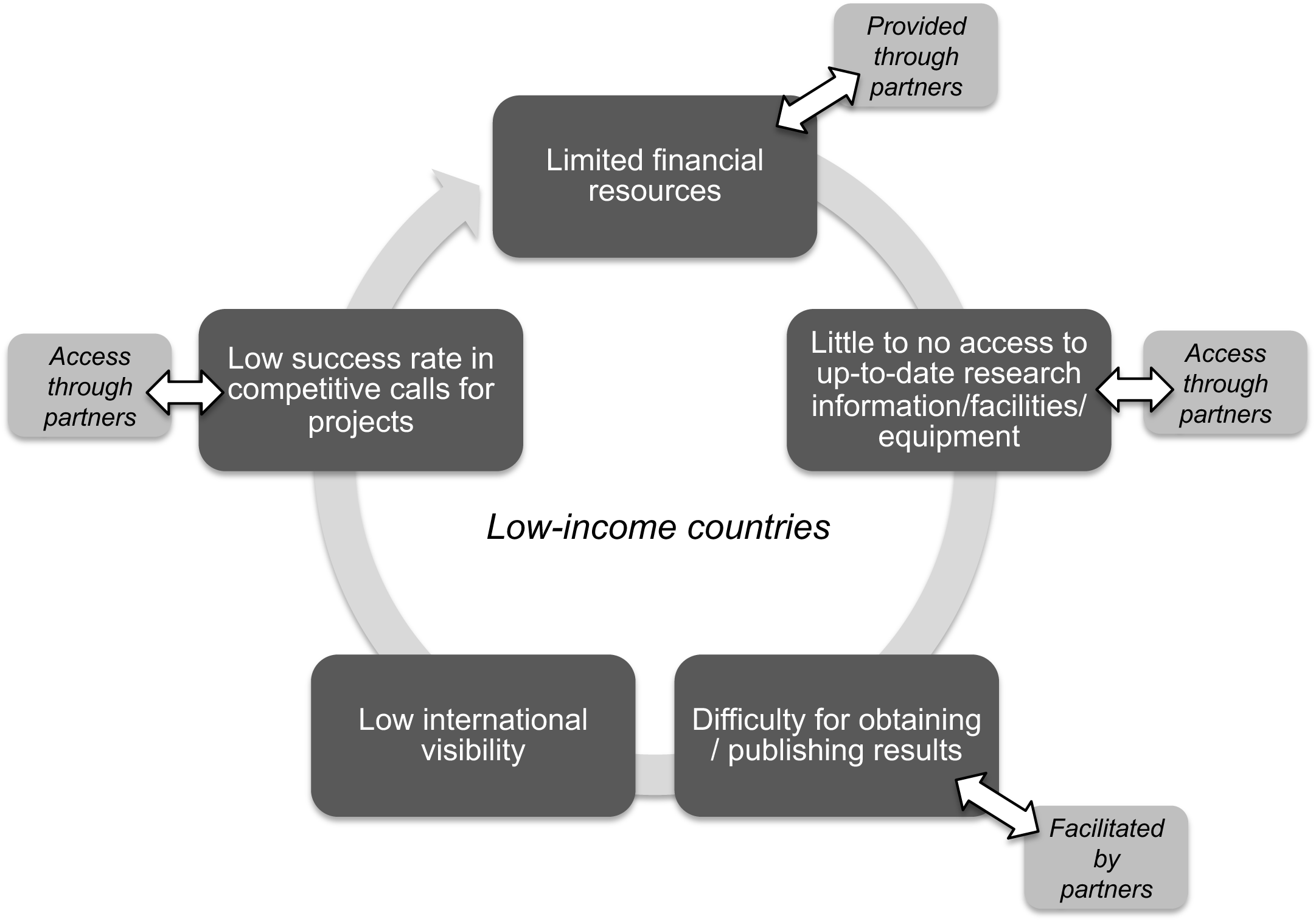
The vicious circle of research cooperation between low- and high-income countries. Several step of the circle can be partially compensated through cooperation with partners from high-income countries, while bringing no definite solution to the central issue (*i.e*. lack of resources) and creating dependency to the collaboration itself.

In this context a change of paradigm is clearly necessary so that West African scientists and teachers may engage into a virtuous circle leading to their self-reliance and the promotion of an upgrade in the research and higher education landscape. Various capacity building interventions working in that general direction have been described previously but to our knowledge, they have mainly targeted specific domains such as Health research, such as CARTA [17] and H3Africa [18,19], and prominently involved English-speaking African countries. We therefore conceived “MooSciTIC: a shot of science!” as a small-scale capacity building project aimed at training the trainers, namely West African teaching assistants, early-career lecturers and research scientists, in cross-disciplinary aspects of their trade. By doing so, we sought to maximize the long-term impact of our training: indeed, we estimated that each teacher from a typical training cohort of approximately 20 trainees would be in charge of teaching to an average of 40 Master students and an even greater number of undergraduate students each year in their respective institutions. Throughout the three years of the project, we could therefore expect to indirectly reach hundreds of students throughout the West African sub-region, with further amplifying effect over the duration of the teacher’s career (Fig 2). Because it is the common practice in the region that a former trainee shares the benefits of their training among their peers within their home institution and sometimes disseminates them further through seminars or workshops, we expected an enhanced impact of the MooSciTIC project.

**Fig 2.**
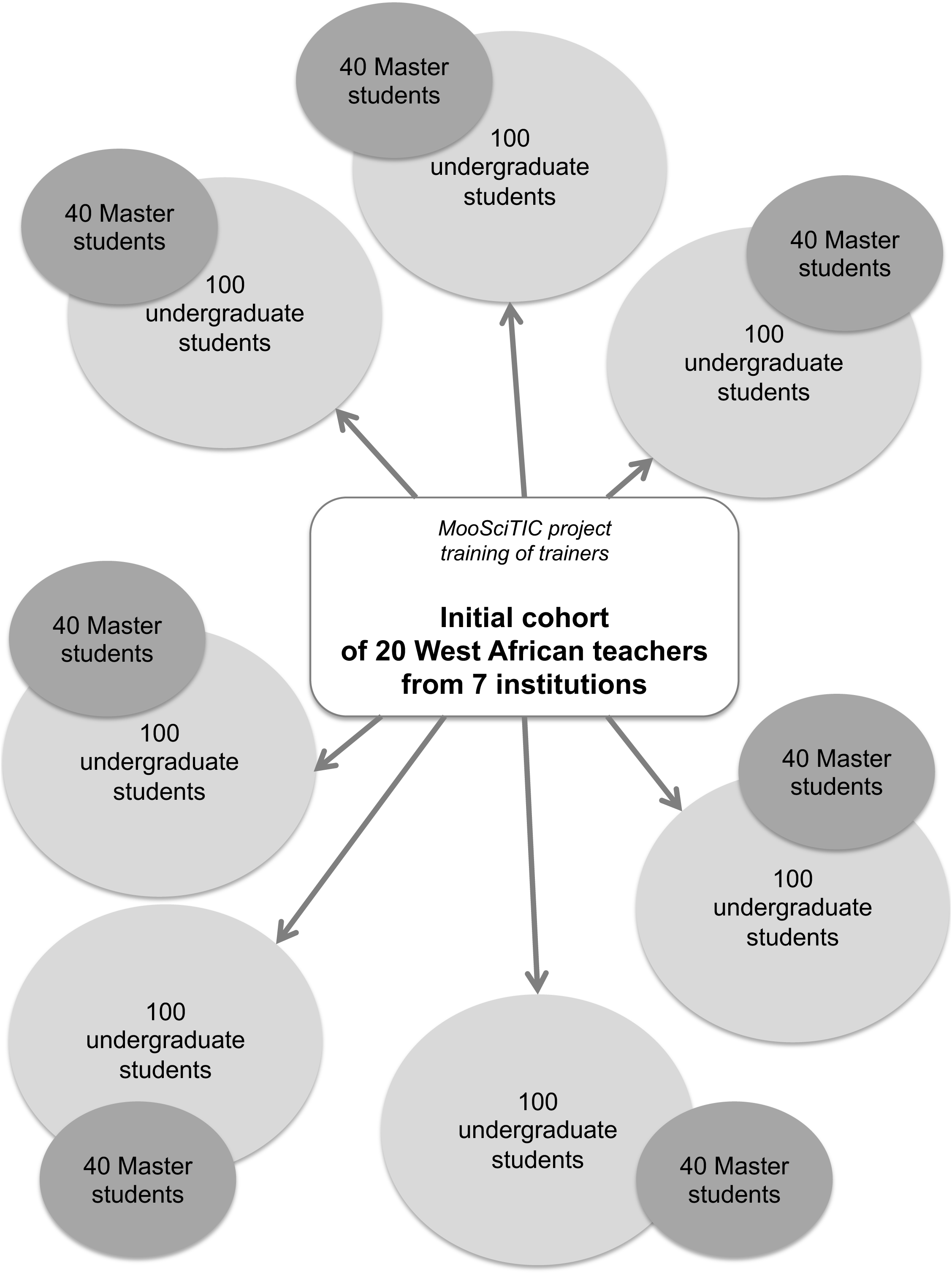
Expected impact of the MooSciTIC project. For clarity, this figure only shows the impact-enhancing effect of the training of a hypothetical cohort of teachers on average numbers of undergraduate and Master students within one academic year.

## Overall structure of the MooSciTIC project

Because we strongly believe that training efficiency is strongly context-dependent, the MooSciTIC project was fully designed, managed and implemented using a collaborative approach involving scientists from France and French-speaking West African countries (who are also the authors of the present paper). Gender parity was ensured in every aspect of the project through joint conception, management and delivery of teaching by male and female scientists and through the active promotion of gender balance among trainees. As illustrated in Fig 3, an initial needs assessment survey was conducted in order to define the training requirements of West African scientists and teachers. The analysis of the responses allowed us to define both the program and contents of the first training session, and to identify criteria for selecting future trainees. Training sessions took the form of a yearly summer school hosted within the University of Abomey-Calavi (UAC), Cotonou, Benin in 2016, 2017 and 2018. For each topic, the contents alternated classical lectures with practical work, mostly by groups of 4-5 people, and left ample time for both group and individual discussions among trainees and between trainees and trainers. The complete teaching materials, software and complementary documentation pertaining to the summer school were provided to trainees on thumb drives. Towards the end of each session, feedback was collected from and discussed with participants in order to help improve future sessions and amend the program accordingly. Outside of the summer school, mentoring was also continuously offered to alumni in order to further help them transpose the benefits of the MooSciTIC training into improved scientific outputs.

**Fig 3.**
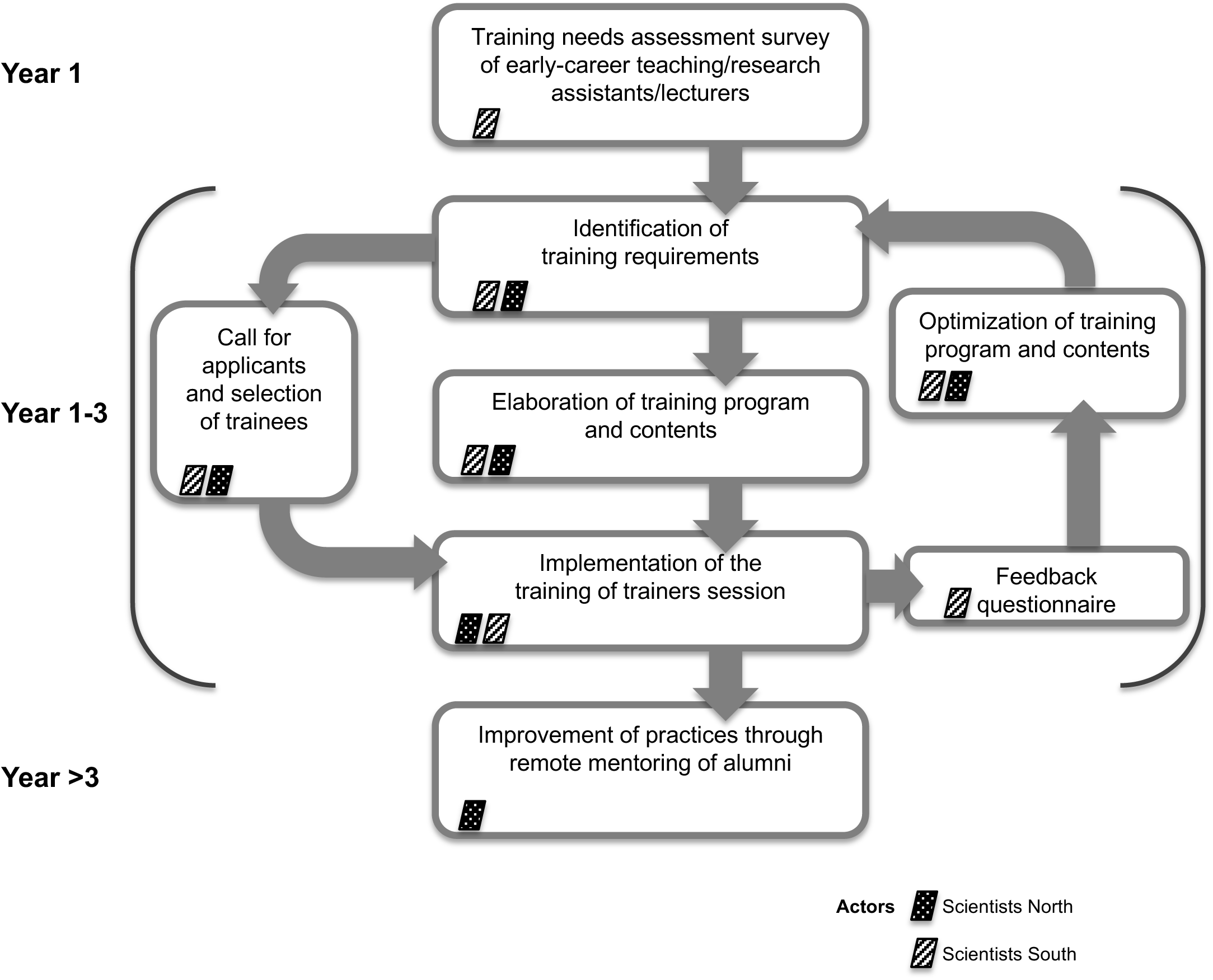
Structure of the MooSciTIC project.

## Training needs assessment survey

At the onset of the MooSciTIC project, we conducted an assessment of the training needs of West African academics in order to validate the rationale of our project. This took the form of a short Google Survey form that was sent by e-mail to contacts and partner institutions throughout French-speaking West and Central African countries. The questions were aimed at assessing the self-perception of the respondents’ skills with respect to either writing scientific articles, writing grant proposals or managing a research project. We received contributions from a total of 19 anonymous respondents from 7 different countries who were involved in undergraduate and postgraduate education in at least 9 institutions (Fig 4A). Data analysis shows that an overwhelming majority of the respondents (from 73.7% for writing papers to 89.4% for writing grant proposals and managing projects; Fig 4B) felt that there was indeed room for improvement in each of these three categories. Among those, a substantial fraction further admitted to lacking prior experience and needing to be provided with basic training on the matter, and represented 27.5% (article writing) up to 58.8% (project management) of the fraction of respondents who had expressed a need for training.

**Fig 4.**
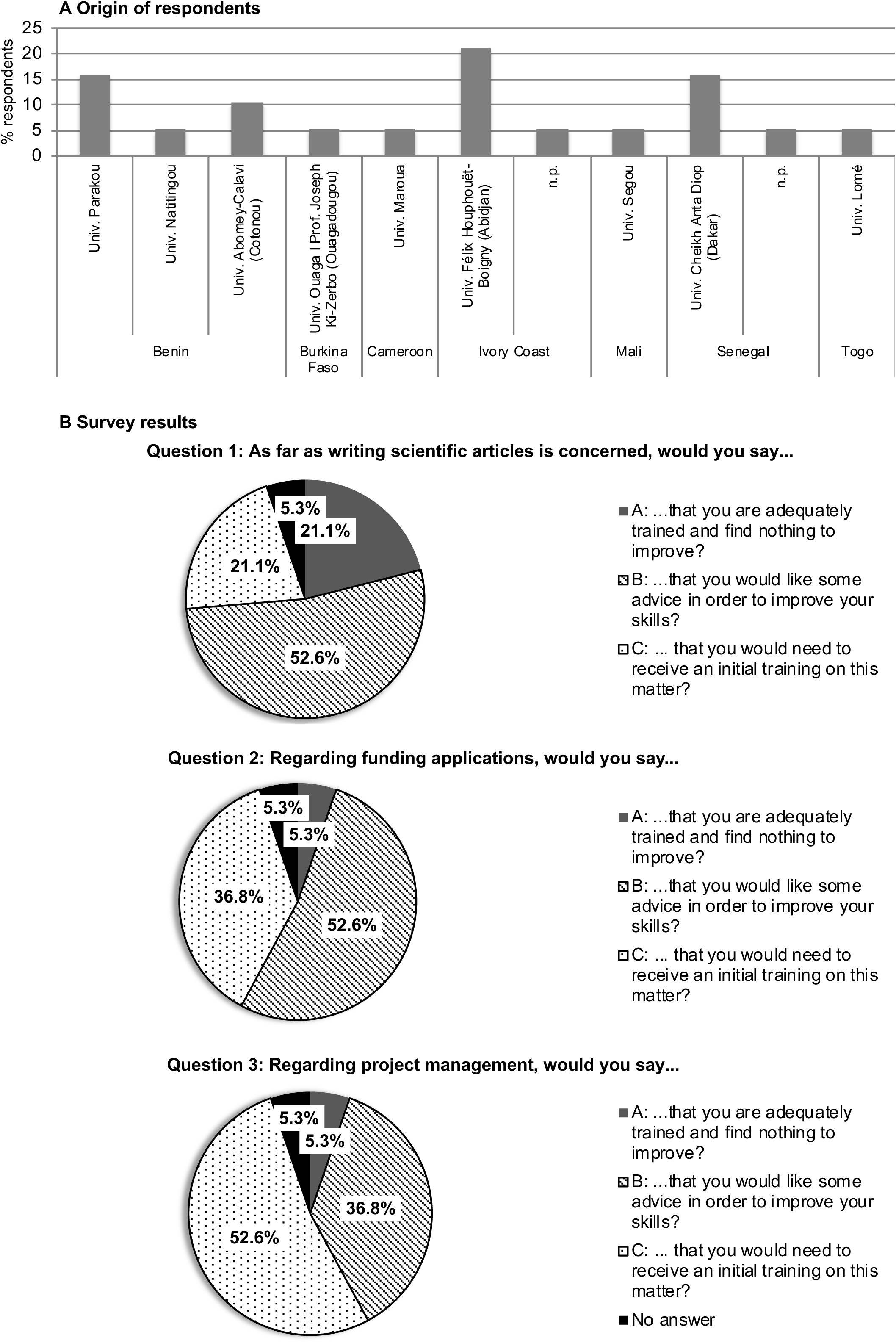
Results of the needs assesment survey. A: Distribution of respondents according to country and institution or origin; B: Distribution of responses to the three questions of the survey. n.p.: not provided.

## Selection process

The call for applicants for the training was advertised by e-mail through both the communication channels used for the survey and West African officed of institutions such as research institutes CIRAD and IRD, and the AUF (Agence Universitaire de la Francophonie; Francophone Universities Agency). In years 2 and 3 of the project, this message was also sent directly to previously unsuccessful applicants. The announcement included a brief overview of the program as well as a hyperlink to an online application form (available here: http://mooscitic.ird.fr/dokuwiki/doku.php) in which applicants were required to provide basic information such as name, e-mail address, gender, institution and country of origin, current position and number of years of experience as a teacher. They were also requested to write a short text explaining their motivation for attending the summer school and upload a 1-page CV. Submission of the application was contigent upon prospective participant’s commitment (materialized by selecting “yes” in a drop-down menu) to re-use at least part of the teaching material within a 2-year period at their home institution. The application process was allowed to run for up to 7 weeks, with 2-3 reminder e-mails sent in the meantime, in order to achieve the desirable number and diversity of applications. Once closed, applications were examined and ranked by a selection committee composed of 2 French and 2 Beninese teachers involved in the project.

In order for the MooSciTIC project to achieve the desired amplifying effect, one of our chief selection criteria was to select participants from as many different countries and institutions as possible. Additionally, in the perspective of inducing emulation through the sharing of experience among trainees, we tried to select participants with different levels of research/teaching experience, and with variable degrees of prior exposure to international research. Since all the teachers involved in the MooSciTIC initiative work in Life Sciences, we chose to select applicants evolving within the same area so that they could relate more easily to our courses.

Ensuring optimal participation from female scientists and higher education staff proved to be challenging. Indeed, it is well documented that because of the stronger social and cultural constraints, women scientists face greater obstacles than their male counterparts for pursuing scientific careers [3,20]. In Sub-Saharan Africa, women represent barely an average 24% of Full-Time Equivalent positions in government-funded agricultural R&D workforce and are less likely than men to reach advanced university degrees (21% have PhDs) or to hold management and decision-making positions (14%) [21,22]. Both formal and informal networks, which are important for accessing opportunities (collaborative research programs, funding, publications and promotions), are predominantly male and therefore poorly accessible to them because of gender discrimination. Despite the fact that we expressly advertized that applications from female scientists would receive special attention, they amounted to exactly a third (81/243) of total applications throughout the project, leaving us no other choice than to use positive discrimination in our selection. Applicants were thus first sorted by gender and origin (country/institution), then ranked within each category according to the relevance of the training to their career path, which led to an initial list of selected applicants who were then contacted by e-mail. As shown in Fig 5, the diversity of each cohort of selected applicants (medium grey) throughout the three years of the MooSciTIC project is a pretty good reflection of the origin and gender distribution amongst initial applicants (dark grey).

**Fig 5.**
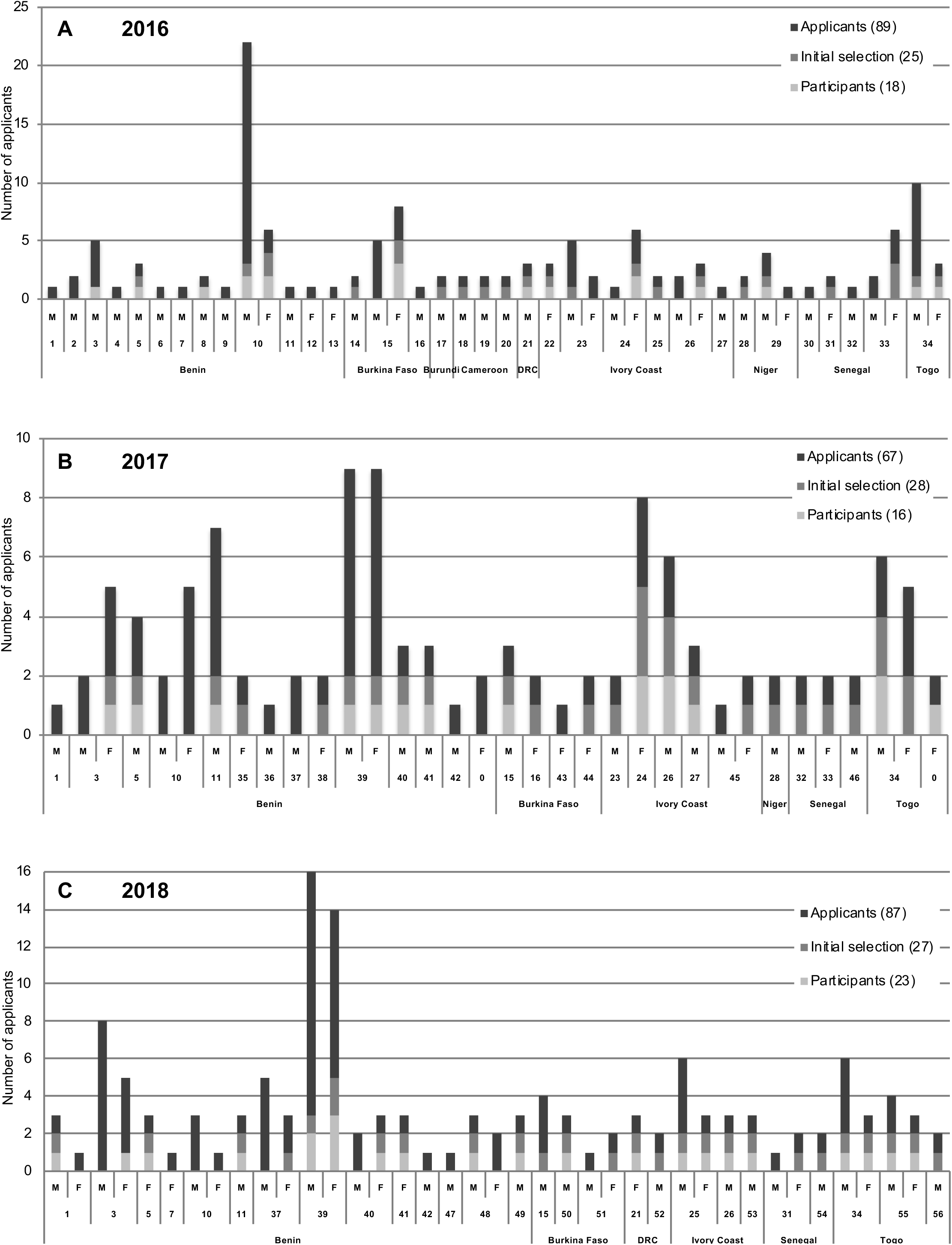
Geographical, institutional and gender diversity among total applicants, selected applicants and participants to the MooSciTIC training sessions. Data are displayed separately for each of the three years of the project: 2016 (A), 2017 (B) and 2018 (C). Selected applicants were included in the initial selection list, whereas participants correspond to selected applicants minus withdrawals plus replacements from the waiting list. M, F indicate male and female applicants, respectively. DRC: Democratic Republic of the Congo. Higher education and research institutions are numbered as follows: 1: Centre Inter-Facultaire de Formation et de Recherche en Environnement pour le développement Durable (CIFRED, UAC); 2: Ecole Nationale Supérieure des Sciences et Techniques Agronomiques, Djougou; 3: Ecole Polytechnique d’Abomey-Calavi; 4: Ecole Nationale d’Economie Appliquée et de Management (ENEAM, UAC); 5: Faculté des Sciences et Techniques (FAST), Dassa-Zoumè; 6: Institut CERCO (private university); 7: Institut National de l’Eau (INE, UAC); 8: Institut Régional de Santé Publique Comlan Alfred Quenum, Ouidah (UAC/WHO); 9: Commission Nationale du Développement Durable; 10: Univ. Abomey-Calavi (UAC), Cotonou; 11: Univ. Parakou; 12: AfricaRice (CGIAR Consortium Research Center); 13: Institut National De La Jeunesse De L’éducation Physique Et Du Sport (INJEPS), Univ. Porto-Novo; 14: Institut du Développement Rural (Univ. Polytechnique Bobo-Dioulasso); 15: Univ. Ouaga I Prof. Joseph Ki-Zerbo, Ouagadougou; 16: Univ. Polytechnique Bobo-Dioulasso; 17: Faculté d’Agronomie et de Bioingénierie, Univ. Burundi; 18: École Nationale Supérieure des Sciences Agro Industrielles, Univ. Ngaoundéré; 19: Univ. Bamenda; 20: Univ. Yaoundé; 21: Univ. Kinshasa (UNIKIN); 22: Centre de Recherches Oceanologiques (CRO), Abidjan; 23: Institut National Polytechnique Félix Houphouët-Boigny (INP-HP), Yamoussoukro; 24: Univ. Félix Houphouët-Boigny; 25: Univ. Jean Lorougnon Guédé, Daloa; 26: Univ. Nangui Abrogoua, Abidjan; 27: Univ. Péléforo Gon Coulibaly, Korhogo; 28: Univ. Tahoua; 29: Univ. Tillabéri; 30: Ecole Supérieure de Génie Industriel et Biologique, Dakar; 31: Institut Sénégalais de Recherche Agricole (ISRA), Dakar; 32: Univ. Assane Seck, Ziguinchor; 33: Univ. Cheikh Anta Diop (UCAD), Dakar; 34: Univ. Lomé; 35: African School of Economics; 36: Centre de Recherche sur le Paludisme Associé à la Grossesse et à l’Enfance (CERPAGE); 37: Faculté des Sciences Agronomiques (FSA, UAC); 38: Faculté des Sciences de la Santé (FSS, UAC); 39: Faculté des Sciences et Techniques (FAST, UAC); 40: Université Nationale des Sciences, Technologies, Ingénierie et Mathématiques (UNSTIM), Abomey; 41: Institut National de la Recherche Agronomique du Bénin, Centre de Recherches Agricoles - Plantes Pérennes (INRAB-CRAPP); 42: Université Nationale d’Agriculture, Kétou; 43: Institut de recherche en sciences appliquées et technologies; 44: Univ. Koudougou; 45: Univ. Alassane Ouattara; 46: Centre d’Etude Régional pour l’Amélioration de l’Adaptation à la Sécheresse (CERAAS); 47: Ecole Normale Supérieure (ENS) de Natitingou; 48: Ecole supérieure Le Faucon, Abomey-Calavi; 49: Univ. Inter Régionale du Génie Industriel des Biotechnologies et Sciences Appliquées (IRGIB) - Africa; 50: Centre International de Recherche-Développement sur l’Elevage en zone Subhumide (CIRDES); 51: Univ. Fada N’Gourma; 52: Univ. Gbadolite (UNIGBA); 53: Univ. Man (U-Man); 54: Institut Supérieur de Formation Agricole et Rurale (ISFAR), Univ. Thiès; 55: Ecole Supérieure des Techniques Biologiques et Alimentaires (ESTBA), Univ. Lomé; 56: Institut Togolais de Recherche Agronomique (ITRA), Lomé; 0: no affiliation. Note that affiliations to different sub-structures within the same complex institution (*e.g*.: UAC) are indicated whenever possible depending on the information provided by applicants.

However, not all selected applicants were finally able to attend, mainly due to either lack of funding for travel and accommodation (see further below) or scheduling conflicts with other professional commitments. The former cause likely played a decisive part in the withdrawal of applicants originating from the most distant countries (Cameroon, Senegal), whereas the latter mostly impacted junior faculty members involved in exam supervision. Also, both the 2016 and the 2017 editions of the summer school overlapped with the Ramadan fasting period, a fact that may have had a negative impact on the participation of Muslim applicants. In any case, we sought replacement for these early withdrawals on a one-for-one basis according to our ranking system. Because of this combination of specific constraints and the strongest emphasis on gender balance, the diversity among the final participants (Fig 5, light grey) was depleted of several institutional and geographical origins compared to the pool of initial applications.

## Design and implementation of the training program

The MooSciTIC project is, by design, aimed at providing its trainees with highly practical information based on recognized standards for good practices and, most importantly, on the sharing of extensive first-hand experience. Indeed, the topics that are at the core of our teaching, such as scientific writing and research integrity, are often not formally taught to scientists (including in developed countries) through formalized courses and practicals, which leaves them to develop their skills through time-consuming trial and errors [23]. In the best-case scenario, mentoring from more experienced colleagues is possible but this option is not always available in institutions from low-income countries and this results in wide variation in research quality [24]. Because of the emphasis on the instant re-usability of knowledge by the trainees in both their daily work and dissemination to peers and students, we used active (participative and collaborative) learning methods since their higher efficiency in that respect has been demonstrated [25,26]. Each topic of the program was thus addressed through sections combining traditional lectures and group/class activities in order to favor the acquisition of on-the-job experience (Table 1). For instance, in the case of the sequence relative to literature mining and bibliographic database management, we designed a course in which the basic principles of a literature search were provided through definitions and examples, completed by a brief overview of search engines and freeware of popular use for bibliographic references management. The latter software were then installed from a thumb drive and set up by the participants on their respective laptop computers. The practical training that followed includes a step-by-step demonstration of the main functionalities of each tool.

**Table 1:** Capacity-building components of the MooSciTiC summer school.

| Topic | Contents |
| --- | --- |
| **Literature mining and reference database management** | *Theory:*<br>• Metadata organization and use;<br>• Principles of web-based literature search: keywords, Boolean operators, wildcards;<br>• Description of search engines (Google Scholar, PubCrawler).<br>*Practice/interactivity:*<br>• Installation of Zotero and Mendeley;<br>• Tour of the main functionalities (importing; searching; formatting citations and lists). |
| **Developing, funding and managing a project** | *Theory:*<br>• Organizing ideas and formalizing a project (mind mapping; concept mapping);<br>• Answering a call for projects (deciphering terms of reference and eligibility criteria; developing the scientific, temporal and financial aspects of the project; anticipating funders' and reviewers' expectations);<br>• Project management.<br>*Practice/interactivity:*<br>• Installation and demonstration of mind/concept mapping tools (Freeplane);<br>• Demonstration of project/task management tools (Cmap, Redmine, Trello);<br>• Responding to a fictional call for projects (group activity);<br>• Individual feedback on a personal draft project. |
| **Scientific communication** | *Theory:*<br>• Basic principles for writing an article (building the backbone: material and methods, figures and tables; trimming experimental data to find the central message; building up the discussion and the introduction in parallel; appropriate referencing; navigating the editorial process and anticipating evaluation criteria);<br>• Tips for improving oral presentation skills and slideshows;<br>• Principles for making an efficient poster presentation;<br>• The scientific CV for grant and job applications.<br>*Practice/interactivity:*<br>• Individual feedback on personal draft articles and posters; |
|  | • Oral presentation of fictional projects. |
| **Mechanisms of scientific investigation** | *Practice/interactivity:*<br>• Case-based activity "An inexplicable disease". |
| **Scientific integrity and research ethics** | *Theory:*<br>• Basic concepts and principles of research integrity;<br>• Illustration through real-life cases involving frauds, misconduct and/or breaches of ethics by scientist;<br>• Ethics in publishing (the case of predatory publishers) and reviewing.<br>*Practice/interactivity:*<br>• Testimonials and debate around personal integrity and ethical issues. |
| **Societal concerns in research** | *Practice/interactivity:*<br>• Testimonials and debate around a societal question of interest for research. |

As for sequences focused on improving scientific writing skills for either articles or grant proposals, each began with a course recapitulating cardinal rules that need to be observed in the presentation of each type of document. Indeed, a large proportion of rejections of submitted manuscripts or grant proposals may be attributed to insufficient attention paid to basic requirements such as suitability with respect to scope and formatting of the journal or call and compliance with writing and organization guidelines, regardless of the scientific quality of the contents [27,28]. Tips for improving writing efficiency and avoiding common pitfalls in the reviewing process were provided, based on both the personal experience of the teachers and several popular “how-to” writing manuals [29–32]. As a complement to these courses, we provided trainees with opportunities to put these principles into practice. Immediately after the course on grant proposal writing, participants were given the text of a fictional call for projects on a somewhat broad theme in Life Sciences, including several options for both budget size and eligibility of different categories of expenses. Trainees were then required to form small work groups and, across dedicated time slots at the end of each working day, each group would build up a grant proposal fitting within the constraints of the chosen project format and the scientific and financial scope of the call. The end-result, composed of a one-page abstract, a financial table and a Gantt chart, was presented to the whole class on the last day of the training session, with ample time allowed for discussions and cross-evaluation among peers. In order to better prepare trainees for this presentation and improve their general presentation skills, a course on principles of good oral presentations was provided beforehand. In parallel, individual mentoring was offered by the teachers in relationship with the article manuscripts and projects drafts shared in confidence by the participants.

So-called “gamified” teaching techniques involving role-playing have been shown to enable easier and longer-term assimilation of complex notions through their first-hand experience in wide a variety of disciplines and learning contexts (see recent examples in [33–37]). During the set-up of the first summer school we stumbled upon the paper from Justin Hines and collaborators [38] describing their use of a simulation sequence built upon real-life events for teaching the basic principles of scientific investigation to students. Because of its elegant simplicity and versatility, we chose to include this activity, named “An inexplicable disease” (see outline in Table 2), to our program, since it might be interesting to our trainees both in itself and as a new teaching tool they might thereafter use with their own students. Briefly, trainees are made to retrace the steps of the scientists who discovered the Kuru degenerative disease in Papua New Guinea in the late 1950’s. The objective is to determine the nature of the disease as either genetic, infectious or environmental, and these concepts are clearly defined through examples at the onset of the activity. Obviously, the actual objective of this sequence is not to solve a puzzle, and indeed most interpretations that tend to emerge spontaneously from the successive experiments are in apparent contradiction with one another. Rather, this exercise is about understanding through experience that scientific investigation is not a linear process and that trials and errors are an integral part of discovery: not all experimental results are readily interpretable in the context of current knowledge, or make sense with respect to the initial hypothesis. Besides, the scenario is designed so that no single group of physicians or anthropologists should be able to reach valid conclusions on their own, and therefore only networking - a process that is essential to research in real life - may steer trainees closer to the correct answer. As a significant bonus, we discovered through implementing this activity that it was a great way to reveal both group dynamics and individual characters, which naturally led us to use it (with great success) on the first day of the 2018 summer school, as an “ice-breaker”.

**Table 2:** Summary of the activity “An inexplicable disease”.

| Objective | Contents |
| --- | --- |
| **Introduction of the activity.** | The instructor gives a brief lecture providing basic background information on the case as well as definitions for the different types of diseases. |
| **Setting up groups.** | Groups of either "Physicians" or "Anthropologists" (*ca.* 4 people/group) are formed. Additional information (different for each specialty) are provided. |
| **Scientific investigation.** | One experimentation (worth 6 to 12 months) is selected by group consensus from a pre-determined list, allowing the group to retrieve new information from the instructor. New experiments may be attempted until the allocated time (24 months) runs up. |
| **Fictional scientific congress.** | A speaker from each group presents the group's conclusions to the class. Among-group differences in both reasoning and results emerge. Speakers navigate between groups to understand the reasons underlying these differences. |
| **General discussion.** | Groups are requested to either amend or confirm their initial conclusions and to justify their decision to the class. |
| **Revelations.** | The instructor gives a lecture on the resolution of the real-life case. |

Scientific research has a strong impact on society through the conceptual frontiers it explores and the innovation it generates. But with great powers come great responsibilities, and, in our highly connected times more than ever, scientists can be held accountable by governments, institutions, private companies and the global tax-paying public alike. This accountability encompasses both what researchers do with the allocated funds, and whether spendings are in agreement with the initial budget items, and (perhaps more importantly) how they do it, *i.e.* whether they conduct their research according to current standards for rigor, traceability and ethics. With this in mind, we wanted to use the MooSciTIC summer school to give our West African colleagues the opportunity to have a glimpse of and take part to current discussions, and to reflect on their own practices and their interactions with society.

This inclusion of societal concerns regarding science and research took two forms. First, as part of each session we organized a debate with local guests to provide testimonies from their own experience. Because of our strong commitment to gender equity, in 2016 we elected “African women in research” as the subject. In 2017, the progressive enforcement of the Nagoya Protocol on Access and Benefit Sharing (ABS) [39] led us to include presentations from the local competent authorities and focal point followed by debate, in a bid to raise awareness on both the additional obligations it induces for scientists and how it entails a new balance of powers in North-South research collaborations.

Our second societal focus was dictated by the increasing public attention gathered by news stories of scientific misconduct or fraud. Over the last few years, several high-profile cases involving scientists of international stature percolated from scientific journals to general news outlets and government officials because of their potential to bear long-term negative consequences on whole research fields through irreproducibility and beyond, on the perception of the trustworthiness of science and scientists by the public and policy-makers [40–42]. Among other outcomes, these recent scandals led to a global realization that awareness of and training to research integrity were insufficient and needed upgrading across most institutions, irrespective of their location in high- or low-income countries [43–45]. The unraveling of one of these cases during the early stages of the preparation of the first summer school inspired us to use it as a counter-example in a case-study based course on integrity and the consequences of fraud. Over the years, the course was gradually improved by adding a larger range of situations illustrating either good or questionable to bad research practices so that the course would not solely be supported by spectacular headline-grabbing stories of deliberate frauds. A greater emphasis was put on examples the trainees could more easily relate to, through the inclusion of commonly encountered cases of unintentional plagiarism and self-plagiarism. It would then open the way to many questions from trainees and lead to a class-wide debate based on the trainees’ personal experience of scientific misconduct-related issues in their work environment.

In addition to these cross-disciplinary contents, theoretical and practical teaching on a variety of topics related to our respective research fields *i.e.* plant genomics and physiology, phytopathology, statistics, bioinformatics and data analyses, phylogenetics, secondary metabolites and pharmacology were included in the program of the summer school, making up as much as 50% of the contents of the 2016 edition. However, various reasons for not maintaining them emerged as the project progressed, as will be discussed in the next section.

## Feedback from the trainees, part 1: post-session questionnaires

In the last days of each summer school session, we provided a questionnaire consisting mostly of open questions to the participants in order to probe their perception of the training (Table 3). Responses were collected anonymously and compiled on the spot, so that the main conclusions could be shared and discussed with all.

**Table 3:** Post-session feedback questionnaire.

| **Question #** | **Question contents** |
| --- | --- |
| **Q1** | How do you rate the overall clarity and consistency of the training session? |
| **Q2** | Were the contents in line with your expectations? |
| **Q3a** | Which part(s) did you prefer? |
| **Q3b** | Explain why. |
| **Q4a** | Which part(s) did you like the least? |
| **Q4b** | Explain why. |
| **Q5a** | Which aspect(s) do you think you will be able to use immediately after the end of this session? |
| **Q5b** | Which aspect(s) do you think you will be able to use later, once you have obtained additional information or training? |
| **Q5c** | Which aspect(s) do you think you will never use? |
| **Q6a** | Do you think that this summer school will help you improve your practice as a teacher? |
| **Q6b** | If yes, explain which part(s) will be the most helpful to you. |
| **Q7a** | Do you think that this summer school will help you improve your practice as a research scientist? |
| **Q7b** | If yes, explain which part(s) will be the most helpful to you. |
| **Q8** | Were the teachers available enough for answering your questions? |
| **Q9** | Do you have suggestions for improving this training session? |
| **Q10** | After this training session, have you identified new needs that you would like to see addressed in future sessions? |
| **Q11** | Would you be willing to recommend your colleagues to attend the MooSciTIC summer school? |

Graphical representations of responses to questions Q1, Q3a, Q4a, Q5a-c, Q6b and Q7b are shown in Fig 6.

The feedback questionnaire generated an excellent overall response rate since it was completed by 100% of the trainees in both 2016 and 2018 and 87.5% (14/16) in 2017. The global clarity and consistency of the program (Fig 6, Q1) was rated as “good” or “very good/excellent” by most of the respondents (83.3% to 100% depending on the year), with a continuous increase of the latter appreciation throughout the duration of the project (from 22.2 in 2016 to 65.2% in 2018). Additionally, contents were seen as partly to completely in line with the initial expectations of the trainees: “yes” to Q2 for 44.4% of respondents in 2016, 85.7% in 2017 and 95.7% in 2018; “yes” with minor negative comments indicating partial satisfaction for 55.6%, 14.3% and 4.3% in 2016, 2017 and 2018, respectively.

**Fig 6.**
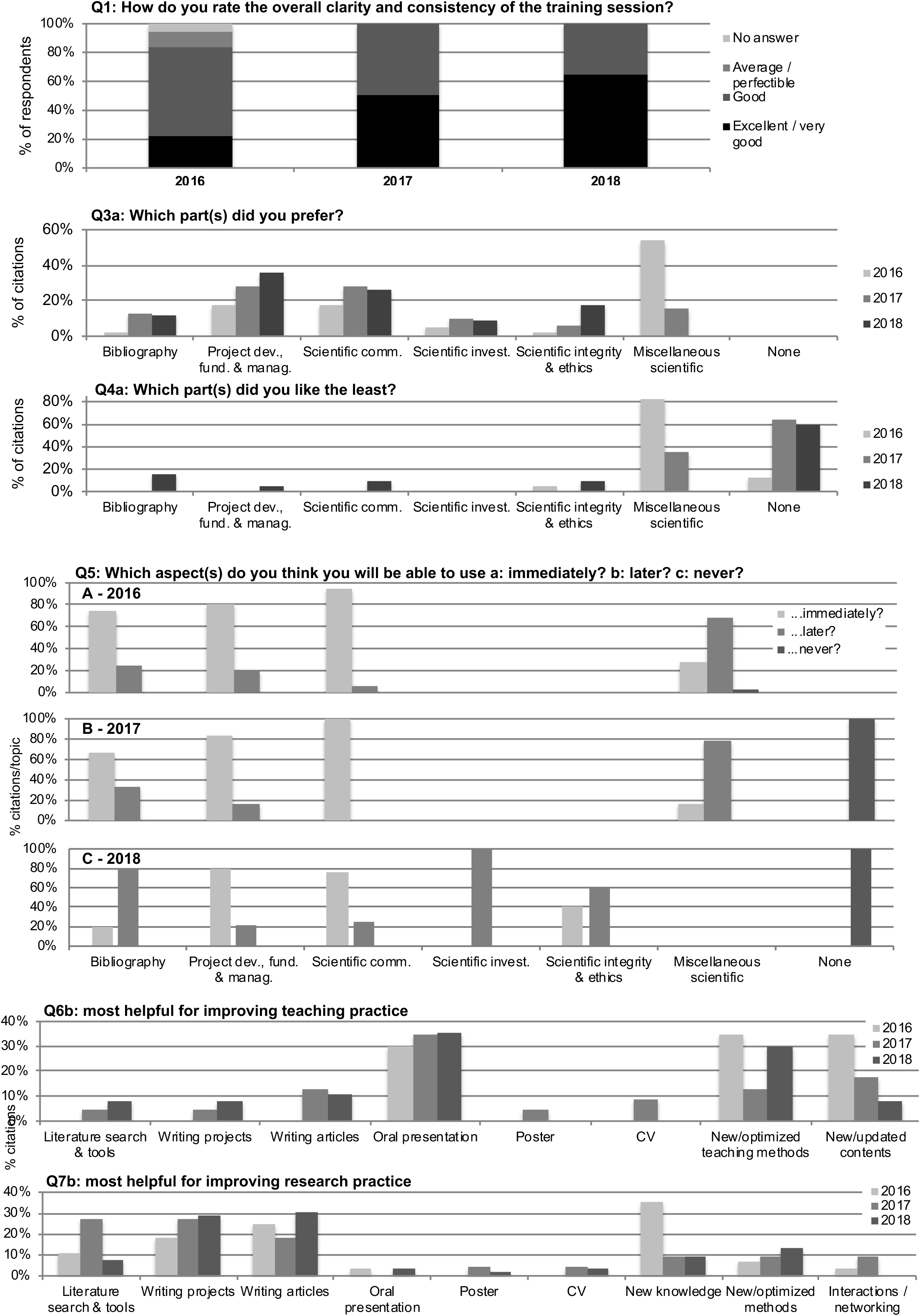
Responses to the post-session feedback questionnaire. See Table 2 for the complete list of questions and text for details.

Throughout the three iterations of the summer school, the aspects of the program that were the most appreciated (Fig 6, Q3a) were those relative to project development and management (from 17.9% of positive citations in 2016 to 36.2% in 2018) and scientific communication (17.9 to 28.1%). The “Miscellaneous” category of teachings including the different science-oriented courses and practicals generated the highest citations rates overall for the 2016 edition, generating both positive (53.8%) and negative (Fig 6, Q4b: 82.6%) perceptions from the respondents. The main reason for dissatisfaction mentioned in the accompanying comments was that insufficient time was allocated for each topic, resulting in poor assimilation, especially from trainees with little to no background. Since the risk of missing the mark because of rising frustration among the trainees seemed to far outweigh the benefits from diversifying the addressed topics, we adapted the program accordingly and reduced science-oriented contents by half. In 2017 satisfaction and dissatisfaction rates for the “Miscellaneous” category had decreased to 15.6% and 35.7%, respectively, however these topics kept on triggering a disproportionate amount of negative feedback compared to other courses. They were therefore completely eliminated from the 2018 session in order to better focus on the cross-disciplinary components of the program.

We then evaluated, sequence by sequence, how soon trainees believed they would be able to re-use what they had just learned (Fig 6, Q5). According to respondents, capacity building topics such as Scientific communication (76-100%), Project development & management (79-83%) and Bibliography (20-75%) had the highest potential for immediate re-use. However, additional training or individual practice was found necessary in 17 to 80% of instances before the latter two could be re-used. In the case of the Bibliography segment during the 2018 summer school, the perception that immediate re-use was unattainable for most participants can be attributed to pervasive issues with the Internet connection that prevented the implementation of the practical that particular year. Strikingly, none of these topics were found of no utility by respondents. By contrast, 69 to 78% of teachings grouped under the “Miscellaneous” category were estimated to be useful only if more training was provided (triggering most responses to Q10), with little difference between 2016 and 2017. This result further confirmed to us that the inclusion of science-oriented topics in the MooSciTIC courses was not efficient as far as the original goal of quickly improving trainees’ self-reliance was concerned.

Overall, 83 to 87% of respondents unambiguously agreed that the summer school would induce improvements in their teaching practice (Q6a). According to them, the most helpful elements for achieving this (Fig 6, Q6b) were the tips for improving oral presentations (30-35% of positive yearly citations), followed by the exposure to new teaching methods and updated, re-usable teaching material. As for the upgrading of their research practice (Q7a), 96-100% of respondents found the session useful, with guidelines for improving project and article writing as well as literature search gathering consistently high citation rates (Fig 6, Q7b: 18-29%, 18-31% and 8-27%, respectively). However, it must be noted that an important proportion of the citations (23-45%) pointed out to more generic aspects as being profitable in the perspective of future research, *i.e.* exposure to new knowledge and new methods and networking opportunities fostered through the training session.

The availability of teachers (Q8) was globally found to be good by 83-93% of respondents. Suggestions for future improvements (Q9) mostly targeted organizational aspects of the training session and attracted mutually incompatible criticisms and propositions. Indeed, daily work schedules were considered to be too heavy and the overall duration of the summer school too long, but it was nevertheless suggested that more time should be dedicated to most topics abnd this would have necessarily implied the implementation of even heavier programs.

Understandably, the emphasis put on international scientific communication at large also underlined how important a good command of the English language is. This point is indeed a major obstacle to the access of French-speaking West African scientist to the international research stage because of an insufficient exposure to and practice of the English language (either in writing or orally) on a daily basis [8]. As a matter of fact, several respondents regretted that we had not included “English as a foreign language” to the program, although it is obvious that no significant improvement in language skills could have been realistically achieved in such a constrained time frame. This demand is also at odds with the fact that, during each session, the use of elements (posters, articles, slides) that had not been translated from English was met with criticisms from some of the trainees.

Although this questionnaire only aimed to assess the perception of the organization and contents of the summer school itself, in many instances issues relative to more logistical aspects were raised by participants. Typically, such comments revolved around the fact that we were not in a position to provide compensations for the travel and lodging expenses incurred by trainees. This was a recurring problem throughout the project since the scarcity of mobility grants sources left most participants with no other option than to self-fund while, in some cases, their salary had been suspended for the duration by their institution. Our decision to gradually decrease the share of science-oriented topics in the program contributed to reduce the duration of the summer school and thereby, indirectly addressed this problem.

## Feedback from the trainees, part 2: “one/two year(s) after” questionnaires

As a complement to the post-session feedback questionnaire, we recently came back to participants from the 2016 and 2017 editions with another set of questions. This time, our goal was to assess the longer-term impacts that the MooSciTIC summer school may have had. We therefore provided them with a Google Survey form listing examples of positive outcomes relative to their research and teaching practice, their career evolution and their networking/collaboration opportunities, and asked them which one(s) they thought might be attributed to benefits derived from MooSciTIC.

Twenty (59%) of the 34 former participants responded and among them, 90% reported improved oral communication and presentation skills, both in scientific meetings and through their courses as teachers; 70% mentioned increased efficiency in the publication of their research work and access to scientific journals of higher quality; 60% reported more efficient student supervision; 40% indicated increased success in competitive grant application processes (remarkably, two respondents reported having obtained more than one grant each). Additionally, 75% of respondents indicated improved networking abilities, mostly through continuing relationships (using e-mails, social media, invitations to workshops and collaborations in research projects) with other participants and teachers. Forty percent mentioned that they had re-used the MooSciTIC teaching material to train students, fellow scientists, or both. Considering the rather low percentages of perceived usefulness of the segment on Scientific integrity and ethics in post-session feedback questionnaires from 2016 and 2017, we were pleasantly surprised to learn that this course was the most frequently disseminated. One respondent even reported having used it as a reference in their contribution to elaborating a code of conduct within their home institution. Finally, 25% of respondents associated their latest promotion to either a tenured position or increased responsibilities to learning outcomes from MooSciTIC.

When asked whether some aspects of MooSciTIC had proved insufficient for achieving the expected improvements, 50% of respondents answered “No” and 35% mentioned the scientific contents. Two of the respondents (10%) were unhappy with their success in grant applications, although one admitted to having worked under an unreasonably short deadline. Finally, one respondent was dissatisfied with the networking opportunities provided, but it was unclear from their answer whether they had actively attempted to keep in touch.

## Outcomes & future directions

By sharing our experience with the MooSciTIC summer school, we aimed at demonstrating the relevance of capacity-building endeavors through the training of trainers and the transforming effect that could be achieved through addressing core aspects in the professional development of research scientists and faculty members. In parallel, we also wanted to show the inner workings of such a training program in the specific context and constraints of West Africa. It was important to us that the latter aspect would include points which, while we had little to no control over them, did have a significant influence on both the implementation of the summer school and its perception and re-use by trainees. We hope that the information included in the present article will inspire other teachers from both the North and the South to engage in capacity-building initiatives at the “training of trainers” level, hopefully with a clearer vision of the opportunities and pitfalls that they entail.

Also, this publication is a way for us to come full circle with respect to our approach to teaching through concrete examples and first-hand experience. Indeed, as part of our teaching material we provided all our trainees with the documents that had been submitted for funding under the MooSciTIC project, as a case in point for successful project development and writing. We see the present paper as another opportunity to provide them with an illustration of how their work and ours, as teachers and scientists, can be promoted and generate added value for the wider scientific community.

True capacity building can only be based on a shift in ownership and leadership of learning contents, results and processes [46,47]. In accordance with this view, it is now in the hands of our West African partners and alumni hands to take over and disseminate the MooSciTIC initiative, while using it for themselves in their daily practice in the field of higher education and research. However, no true and lasting transforming effect can be achieved without the active support from governments, local policy-makers and heads of institutions [4,5]. Their willingness to provide incentives for upgrading the training of teachers and scientists and favor the development of structured, jointly coordinated national and trans-national/regional research networks is key for conquering an independent future for West Africa.

## Acknowledgements

We are grateful to all participants of the MooSciTIC summer schools for insightful discussions, feedback, and their unfailing support to the project. We also thank Anne-Laure Roy (CIRAD) for the critical reading of the manuscript. The work described in this article was funded as part of the MooSciTIC project supported by Agropolis Fondation under the reference ID 1501-011 through the “Investissements d’Avenir” Program (Labex Agro: ANR-10-LABX-0001-01). This is publication ISEM 20xx-xxx.

## Ethics statement

Surveys, application forms and questionnaires included disclaimers stating that collected data would only be used after anonymization and for the purpose of project assessment only. Gender parity was enforced consistently throughout the project, among coordinators, teachers and trainees. All teaching material produced and/or used as part of this project is distributed under the Creative Commons license Attribution - NonCommercial - ShareAlike (BY-NC-SA) 4.0 International (http://creativecommons.org/licenses/by-nc-sa/4.0/) and is available upon request.

Author contributions
Conceptualization: all authors Funding Acquisition: all authors Project Administration: EJ Supervision: LL, EJ Resources: all authors Data analysis and visualization: MA, KA, EJ Writing – Original Draft Preparation: all authors

